# Nanoscale imaging of native symbiotic animal tissue using a multimodal large volume imaging pipeline for cryo-electron tomography

**DOI:** 10.1101/2025.11.30.691379

**Authors:** Katrina A. Gundlach, Oda H. Schiøtz, Mark Ladinsky, Colin Raimann, Michael Rheinberger, Florian Beck, Barış Gündüz, Rob Langelaan, Martin Rücklin, Ronald W.A. Limpens, Edward G. Ruby, Margaret McFall-Ngai, Jürgen M. Plitzko, Ariane Briegel

## Abstract

The field of cryo-EM offers the possibility to gain high-resolution structural information of biomolecules in their native state. Advances in sample thinning of cryo-EM samples allows the study of proteins inside intact cells using tomography, opening the door for ‘visual proteomics’. However, thicker samples such as tissues or entire organisms are still largely unsuitable for cryo-electron tomography (cryo-ET). Therefore, significant efforts are directed toward developing and improving preparation methods to enable cryo-ET of such complex samples. We focused on the binary association between the Hawaiian bobtail squid *Euprymna scolopes* and the luminous bacteria *Vibrio fischeri*. The Squid–Vibrio system has long been studied to understand host-symbiont interactions. Our goal is to study the bacterial-host interface using cryo-ET, at a resolution previously unattainable by conventional EM methods. Here, we present a multi-modal preparation and correlative imaging workflow—including cryo-fluorescence microscopy, microCT, freeze-substitution electron tomography (FS-ET), and serial blockface SEM—to localize and prepare specific regions of the dissected symbiotic light organs for cryo-ET. This approach enabled us to directly visualize symbiotic *V. fischeri* within the internal host crypts at macromolecular resolution, revealing spatial organization, physical contact, and putative exchange interfaces between host and microbe. Our findings provide structural insights into a foundational model of host–microbe symbiosis and demonstrate the feasibility of cryo-ET for investigating intact tissues at the nanoscale.

## Introduction

Host-associated microbes play vital roles in the development, physiology, and health and disease of animals, plants, and fungi. The structural basis of these host-microbe interactions, however, remains essentially unexplored at the nanoscale level despite being essential in understanding the roles of cellular components at the interface between species. For example, many important microbial factors that play a role at this interface, such as the type-6 secretion system of gram-negative bacteria ^1–3^, have not yet been studied within the context of the tissues of an intact host. More broadly, major questions have yet to be answered concerning the interaction between microbes within a host, and the interaction of microbes with the host tissue itself. While light microscopy preserves sample integrity, its resolution is insufficient to visualize these interactions at the nanoscale. Conversely, traditional electron microscopy offers subcellular detail but requires harsh sample preparation that can obscure or distort native architecture. As such, we have chosen *in situ* cryo-electron tomography (cryo-ET) as the best method by which to explore these questions.

Cryo-ET enables imaging of biological samples in a near-native state at macromolecular resolution. Its application, however, has been largely confined to samples with an extremely thin depth like isolated protein complexes, viruses, or small bacterial cells until more recently. Advances in cryo-ET sample preparation techniques, such as the ability to trim vitrified tissue samples using focused ion beam (FIB) milling, has opened the door to imaging larger biological samples including intact tissues and even small organisms with cryo-ET ^4^. More specifically, recent developments in the large-volume sample processing pipeline provided proof of principle that such complex tissues are amenable for cryo-ET^5,6^. Here, we adapted the Serial Lift-Out method, which creates a series of very thin slabs of tissue called lamellae (25 µm x 45 µm x 200 nm) prepared from a larger block of tissue extracted from a large high-pressure frozen specimen (<200 µm thick)^5^.

We applied this cryo-ET Serial-Lift-Out pipeline to study the binary symbiosis between the Hawaiian bobtail squid, *Euprymna scolopes*, and its specific bioluminescent symbiont, *Vibrio fischeri*, a model for animal–microbe interactions. The light-emitting organ in which the bacteria are maintained provides a platform for the study of the colonization of animal epithelial with bacteria, but is much simpler than most other symbiosis-model systems due to the binary relationship between the two organisms ^7^. The Squid—Vibrio model system has been previously studied using a variety of approaches, including metabolomic ^8^, transcriptomic ^9^, computational ^10^, and confocal light microscopy^11–16^. The system also has a long history of conventional transmission electron microscopy that provided the first glimpses into ultrastructure of the system in both juvenile and adult light organs^17–21^. These studies have provided deep insights into host-microbiome interactions. For example, the interactions of bacterial flagella and secretion systems with host tissue have been described in detail at the light microscopy level ^22–25^. Despite these extensive studies, much remains unexplored at the nanoscale, with cryo-ET imaging providing a means to understand the structural aspects of this intricate interaction.

In this study, we developed a multi-modal preparation and correlative imaging workflow—including cryo-fluorescence microscopy, microCT, freeze-substitution electron tomography (FS-ET), and serial blockface SEM—to localize and prepare specific regions of the squid light organ for cryo-ET. Our approach enabled direct visualization of symbiotic *V. fischeri* within the internal host crypts at macromolecular resolution, revealing spatial organization, physical contact, and putative exchange interfaces between host and microbe. These results provide structural insights into a foundational model of host–microbe symbiosis and demonstrate the feasibility of cryo-ET for investigating intact tissues at the nanoscale.

## Results

To answer questions about the structural interactions at the host-microbe interface, we needed to accurately find a very small target within a much larger piece of tissue. The dissected light organ of a juvenile squid has dimensions of ∼700 µm x 150 µm x 400 µm, which corresponds to an approximate volume of 50 million µm^3^. As a comparison, a reconstructed tomogram collected using cryo-ET covers a minute fraction of this volume (1 µm x 1 µm x 200 nm), which corresponds to an approximate volume of 0.2 µm^3^, i.e. 250 million tomograms would fit inside a single light organ. This targeting can be aided by previous work performed with histological sections, conventional EM, and confocal microscopy which has provided the foundation for our current understanding of the architecture of the light organ. This knowledge provided a structural basis for sample orientation and for contextualization of the cryo-ET data. These imaging modalities, however, were insufficient for accurate targeting of host-microbe interface in the cryo-ET samples given that they were performed on separate, often chemically fixed, samples.

To overcome this challenge, correlative approaches using fluorescent labeling are commonly applied to accurately localize the biological target during FIB-milling for cryo-ET, *e.g*., using GFP-tagged symbionts within crypt spaces. However, the fluorescent signals deep within the tissue of large-volume cryo-samples are often weak, and autofluorescence in surrounding tissues, highly prevalent in cryogenically preserved samples, can lead to false targeting^26^. These problems can be partially addressed by fluorescently labeling and screening the entire tissue with a different fluorescent tag such that, even if the area of interest resides deeper within the tissue, the shape of the tissue can be used for targeting.

### MicroCT

To gain insight into the gross 3D architecture of the light organ in the context of the surrounding squid tissue, we applied micro-computed tomography (microCT). This method images objects in 3D using X-ray CT. For this procedure, the light organ needed to be removed from juvenile squid. The tissue was dissected along with some of the adjacent tissue, including the ink gland, ink sac and hindgut, as it cannot otherwise be cleanly separated. The sample was vitrified, followed by freeze substitution (FS), plastic embedding and staining (see Materials and Methods).

Our MicroCT data revealed the entire light organ and connective tissue in a three-dimensional (3D) model that we could spatially freely rotate. The structural features of the dissected tissue complex were easily identifiable, such as the appendages, ink sac, ink gland, hindgut, and symbiont migration path from the surface pores to the inner crypt spaces (Fig. 1A-C). By comparing this model to our cryo-fluorescent data, it became apparent that the ink gland was highly autofluorescent and overpowers the fluorescent signal from the crypt spaces filled with GFP-tagged bacteria. This insight was essential to allow accurate targeting for the subsequent cryo-EM workflow steps. Additionally, the dataset provided a good estimate of expected depth of crypt-space localization relative to the surface of a frozen sample.

**FIG 1.**
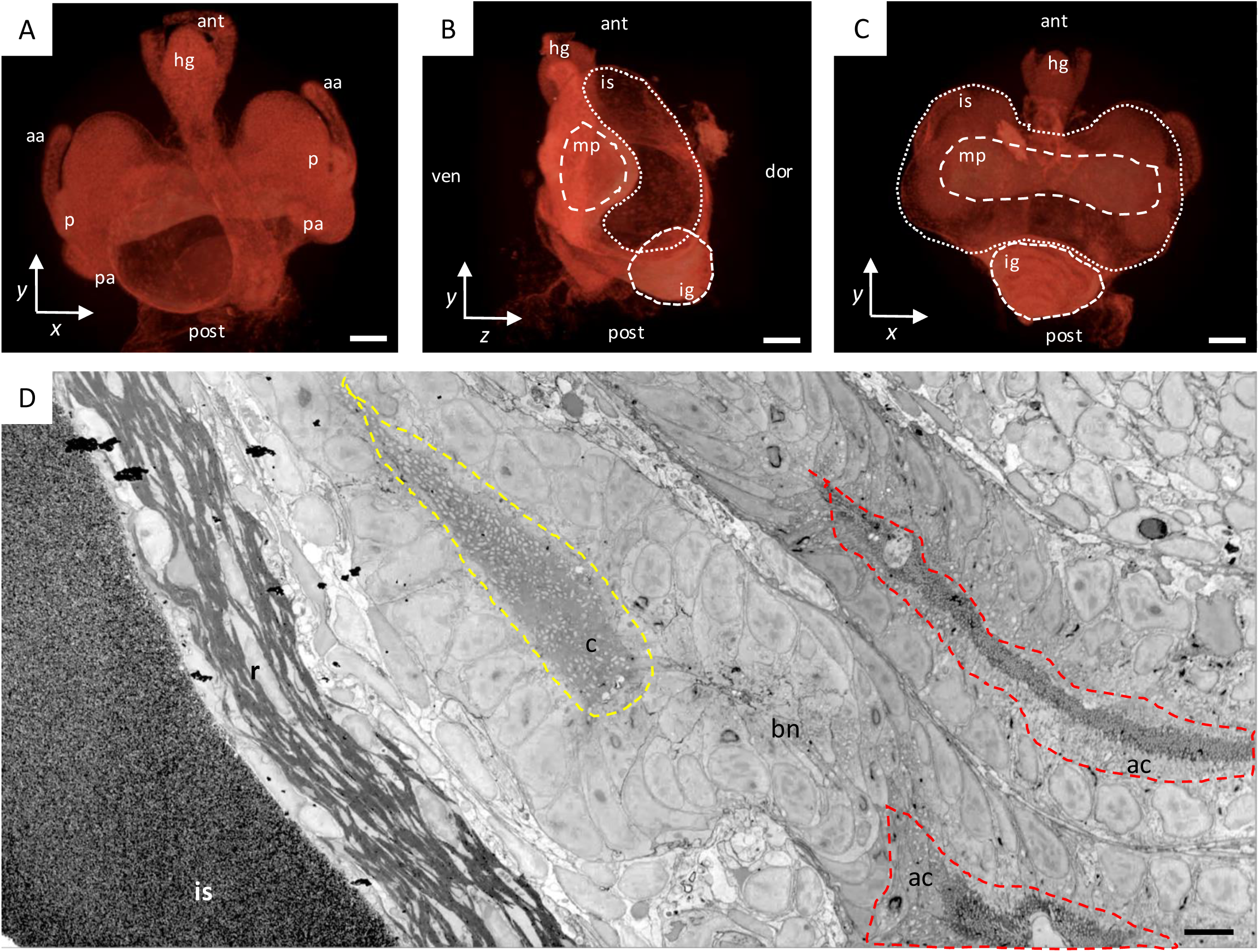
Micro-CT and serial blockface SEM of juvenile *E. scolopes* light organ. A-C) Still images from MicroCT data collection, (aa) anterior appendage, (pa) posterior appendage, (p) pores, (hg) hindgut, (mp) migration path, (is) ink sac, (ig) ink gland, (c) crypt, (bn) bottleneck, (ac) antechamber, (r) reflector, (ant) anterior, (post) posterior, (dor) dorsal, (ven) ventral. Scalebars: (A-C) 50 µm, (D) 10 µm.

**FIG 2.**
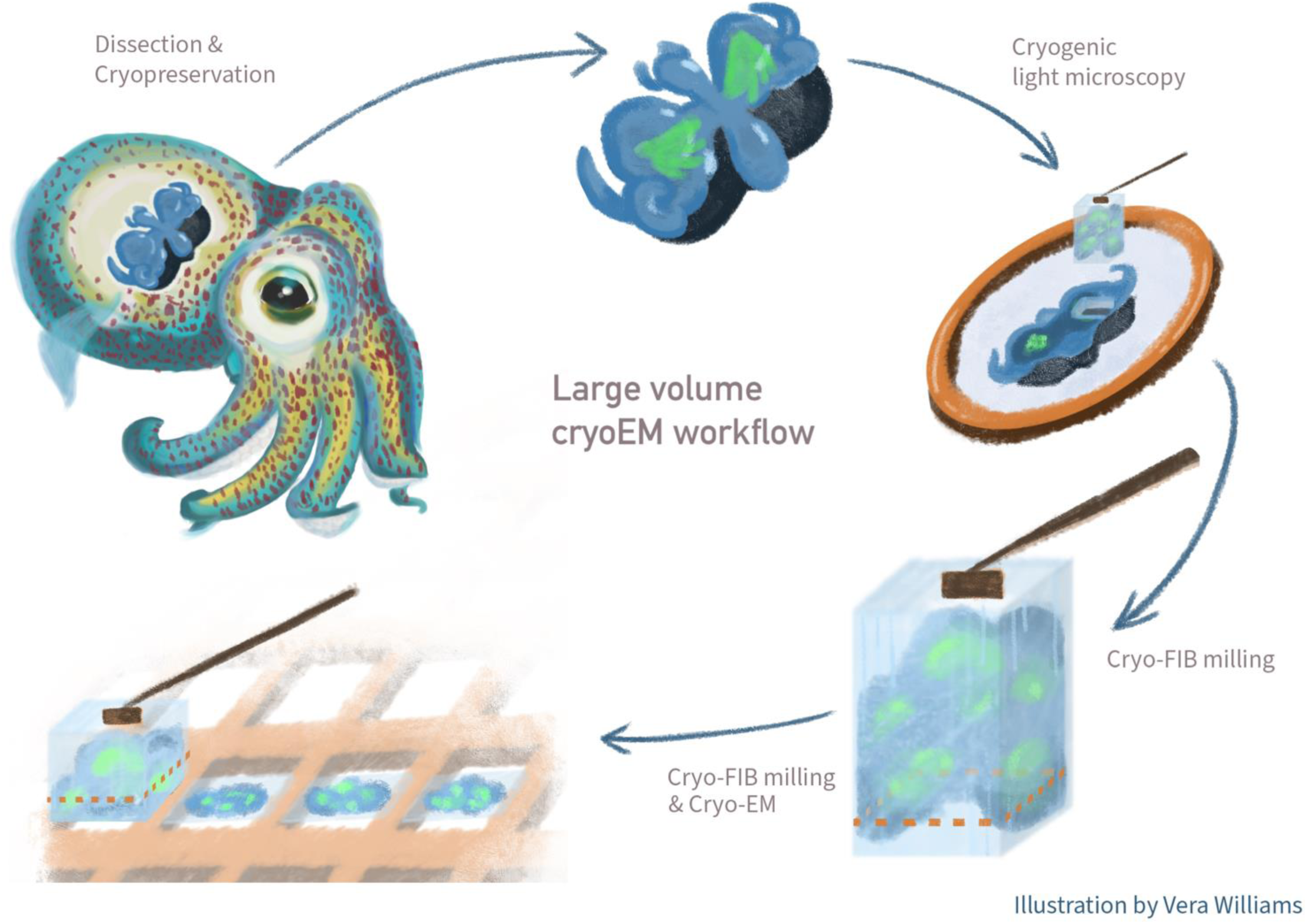
Large volume sample workflow for symbiotic light organ of Hawaiian bobtail squid for cryoET. Light organs colonized with GFP-*Vibrio fischeri* are placed in HPF planchet with cryoprotectant and HPF. Lift-out block of host-microbe interface is removed via cryo-FIB SEM, serially sectioned and attached to grid bars of lift-in EM grid. Prepared lift-in grid is then imaged for cryo-ET.

### Serial Blockface SEM

While MicroCT provided a clear general overview of the organ-tissue complex in 3D, more detailed information at the cellular level, such as insight into the different cell types inside the tissue, was not retained. To obtain these data, we used serial blockface scanning electron microscopy (SEM) to image the crypt space and the surrounding host tissue. This method required the tissue to be processed following established traditional EM preparation workflows, resulting in dehydrated, plastic embedded, and stained tissue samples. For imaging, the sample was then inserted into an SEM, and an integrated microtome iteratively removed the top slice from the sample surface followed by SEM imaging.

Our serial blockface SEM data showed the light organ tissue at cellular resolution directly in and around the crypt space (Fig. 1D). It allowed us to follow the host-microbe interface in three dimensions along the structurally intricate migration paths using the other anatomical features as biological landmarks. Clearly discernable tissues and cell types included the crypt spaces filled with bacteria, the microvilli-lined epithelial cells of the crypt and cilia-lined migration path through the antechamber, neuron/blood sinuses, reflector, lens, and ink sac. This imaging information was used in the final stages of our cryo-EM large volume workflow in order to contextualize the data. Here, the remaining surface area of the lamella was limited to ∼20 µm x 30 µm. This small field of view contained only limited biological context and the distinct landmarks obtained with serial blockface imaging were key in understanding the cryo-ET data.

### Cryo-ET

The ultimate goal of this work was to use *in situ* cryo-ET imaging to investigate the microbe-host interface in a near-native state, at macromolecular resolution. However, a number of challenges exist when preparing such a large volume specimen, one challenge is the need to start with proper vitrification. In contrast to thin-sample preparation where the sample is plunge-frozen directly on an EM grid^27^, thicker samples, exceeding a few µm, need to be prepared using high-pressure freezing. This requirement is due to the cooling rate during plunging being too slow to prevent ice crystal formation which, in turn, destroys the ultrastructure of the sample^28,29^. To vitrify larger specimens, pressure is applied during the rapid cooling process to prevent sample expansion and thus preventing ice crystal formation. Despite the method of freezing and multiple optimization rounds, still not all samples were fully vitrified and only samples that were well vitrified could be used for subsequent data collection.

After optimizing the multimodal imaging steps outlined in the previous section, we were able to successfully target the region of interest in our vitrified light organs using the obtained contextual information and correlative cryo-light microscopy of the host tissue and bacterial GFP tag. This guided the preparation of the block for extraction and section, using the Serial Lift-Out method (Supplementary Fig. 1). This approach ensured that the extracted region in the frozen sample block that was lifted out and serially sectioned contained the desired regions. This outcome was further confirmed by fluorescence imaging of the sections using the integrated LM inside the FIB instrument, where the individual sections clearly showed the expected GFP signal from the fluorescent bacteria (Supplementary Fig. 2A).

### *In situ* host tissue architecture

When exploring the host tissue surrounding the crypt spaces, we observed expected cellular structures, albeit at unprecedented resolution and structural preservation (Supplementary Fig. 2). These features included cell-cell junctions between the epithelial cells and flotillin-like structures at the base of the microvilli (Fig. 3A, C). Additionally, typical biology such as mitochondria, Golgi apparatus, and nuclear pores were observed throughout the host tissue (Fig. 3A, B, and D, respectively). As the material extended past the epithelia, other host material was observed such as muscle cells and extracellular matrix (ECM) (Fig. 3D, E).

**FIG 3.**
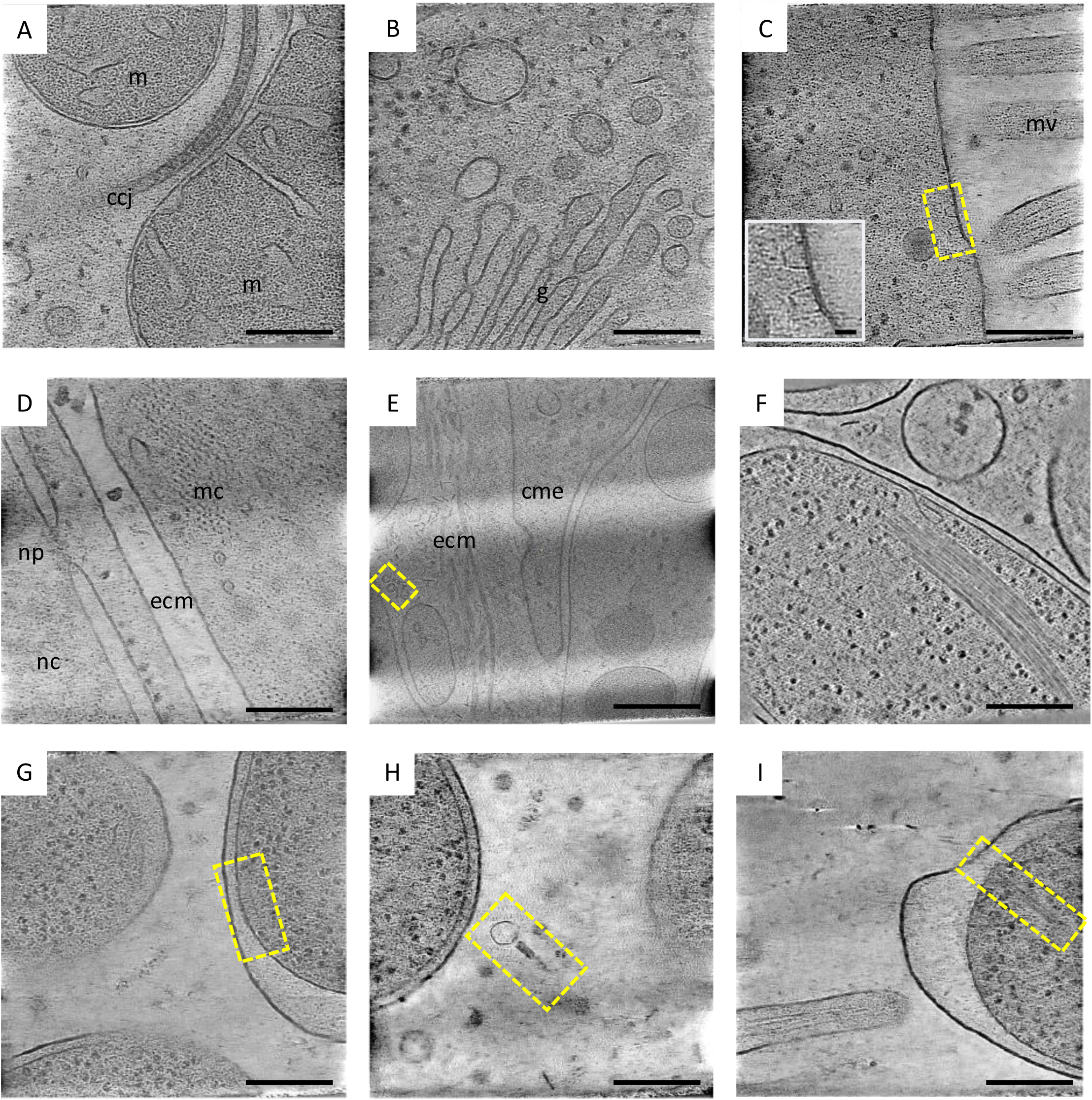
Cryo-ET of host tissue and microbe nanostuctures. (A) Cell-cell junctions (ccj) between epithelial cells with mitochondria (m), possibly resembling adherence junctions. (B) Apical side of epithelial cells near crypt space, showing large areas of Golgi apparatus (g). (C) Flotillin-like structures found at the apical membrane of the host epithelial cells, microvilli (mv). (D) Muscle cell (mc) with actin and myosin filaments near nucleus (nc) and its nuclear pore (np), including extracellular matrix (ecm). (E) Haemocyanin (yellow box) near extracellular matrix (ecm) during clathrin-mediated endocytosis (cme). (F) Bacterium with clear bundle of filaments of unknown identity. (G) Chemotaxis array in yellow box. (H) Bacteriophage within crypt space (yellow box), empty head indicating this phage is post-infection. (I) Type 6 secretion system (yellow box), empty sheath indicating the T6SS is post-discharge. Scale bars (A-D and F-I) 200 nm, (E) 400 nm, (C) inset 20 nm.

Unexpectedly, additionally within the epithelial cells regular crystalline-like arrays of filaments were observed (Fig. 4). They could be clearly observed at lower magnification and were deemed interesting as little is known about large filament arrays in epithelial cells (Fig. 4A). In order to determine the identity of the filaments, the hexagonally packed filaments were averaged to a resolution of 14 Å, GSFSC (Fig. 4B, C, Supplementary Fig. 3A-D). This structure showed a non-polar filament made up of three homodimers rising with a ∼120° twist and with a helical rise of ∼290 nm. When comparing this with previously solved filaments structures, it closely resembles the inactive state of human acetyl-CoA carboxylase (ACC), which has previously been solved in both its active and inactive states^30^ (Supplementary Fig. 3E). The structure differs, however, in that the structure solved here has an additional density resembling a coiled-coil domain that seems to span the distance between the filaments, likely resulting in the maintenance of the crystalline-like arrays. While the identity is not certain and further biochemical assays will be needed to confirm the identity of the filaments, if indeed this is squid ACC, it would be the first time such a filament has been shown to form crystalline-like arrays. Given its resemblance to the inactive form, it would also be of interest to determine whether this crystalline-like array is maintained in the active state of the molecules, or whether it is the dissociation of the crystalline-like lattice that results a conformational change and, in turn, the activation of the molecule.

**FIG 4.**
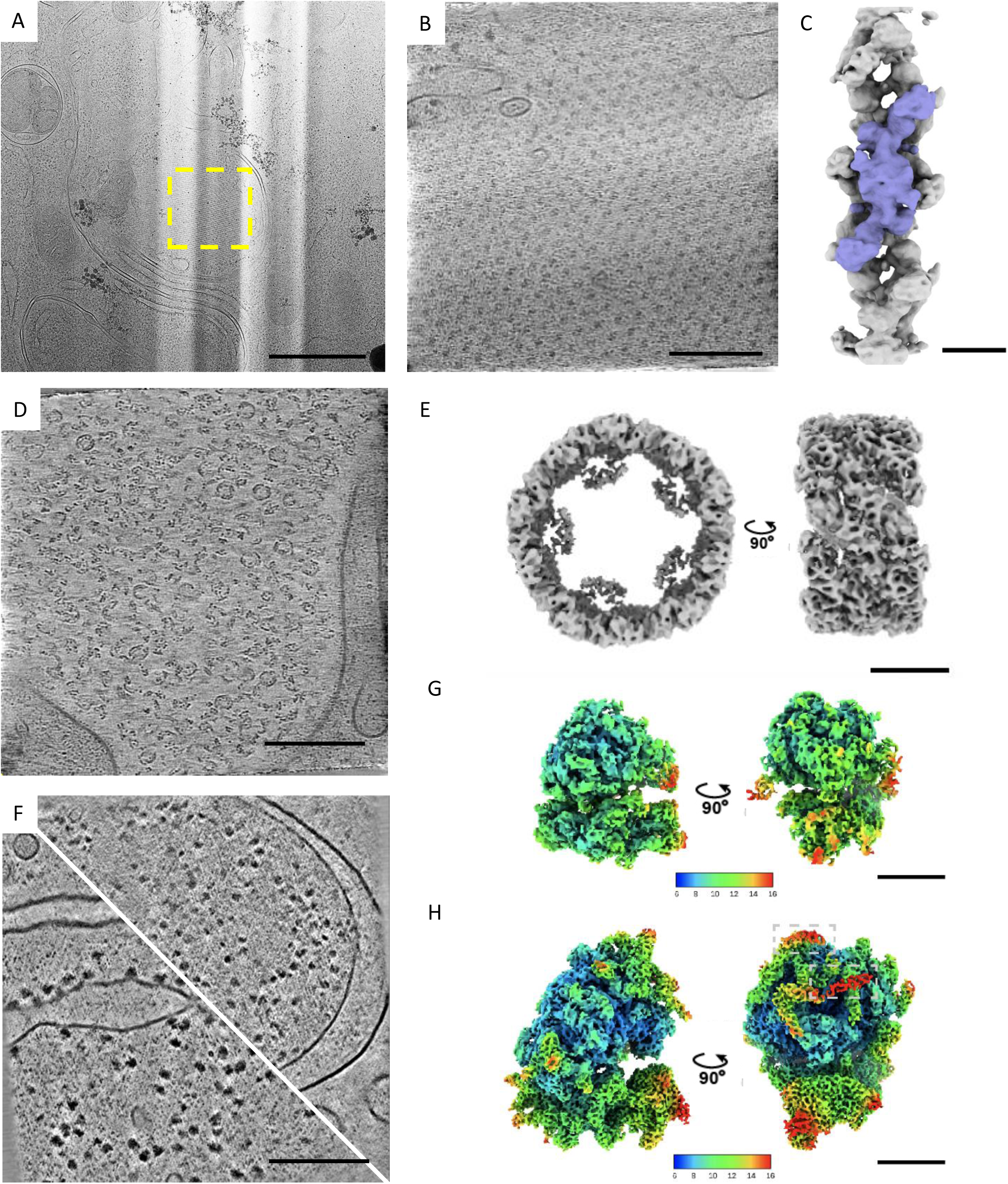
Subtomogram averaging of host and microbial structures. (A) Low magnification (11,500 × magnification) image of an epithelial cell containing the crystalline-like filament array (yellow box). (B) Slice through a tomogram containing the crystalline-like filament array. (C) Structure of the filament resolved to 14.2 Å resolution, GSFSC. Purple highlights the homodimer, three of which make up a full ∼290 Å helical rise. (D) Slice through a tomogram of squid blood sinus filled with haemocyanin proteins. (E) Subtomogram average of haemocyanin protein from squid tissue resolved to 9.9 Å, GSFSC. (F) Slice

Other distinctive features found in the host tissue were haemocyanin complexes, located throughout the animal tissue extracellular matrix (Fig. 3E, Supplementary Fig. 4B) and within blood sinuses (Fig 4D, Supplementary Fig. 4A). Haemocyanin is known to be involved in the onset and maintenance of the symbiosis between *V. fischeri* and the light organ, where one of the main roles is delivering oxygen required for bioluminescence to the crypt spaces. Haemocyanin is distinct in shape, barrel-like from top-view (Fig. 3E, Supplementary Fig. 4C) and having an hourglass-like shape from side-view (Fig. 3E, Supplementary Fig. 4D) and it is seen throughout the animal tissue but, so far, not in the crypt space of juvenile animals. While previous single particle cryo-EM and crystal structures have been reported, we were able to resolve the first *in situ* structure of haemocyanin to a resolution of 9.9 Å, GSFSC, using subtomogram averaging. This structure closely resembled the structure solved from Japanese flying squid, with a D5-symmetric outer collar ^31^. We were, however, unable to clearly resolve the asymmetric inner collar as was done with the single-particle structure.

### Distinction between organisms using ribosome structure comparison

When inspecting the acquired data, the distinction between the host tissue cells and the bacteria was straightforward. However, while this capability was not a challenge here, there was a possibility that accurate cell identification between organisms might not always be unambiguous. Therefore, as a proof of principle, we used subtomogram averaging to differentiate between host and bacterial structures, and we averaged the ribosome structures of both species. While these macromolecules perform the same role in both organisms, they were quite structurally distinct, with the bacterial 70S ribosome being ∼25 nm in height (Fig 3G, Supplementary Fig. 5A), and the squid 80S ribosome having a height of ∼32 nm (Fig 3H, Supplementary Fig. 5C). The latter had a loop of additional density not previously seen in ribosome structures, perhaps unique to the cephalopod lineage. When resolving the two structures, the squid ribosome was refined as a singular structure to an average resolution of 7.5 Å, GSFSC (Supplementary Fig. 5E), while the bacterial ribosome had to have its small and large subunits refined independently to 9.7 Å and 8.7 Å, GSFSC, respectively (Supplementary Fig. 5B, C). The latter is indicative of greater movement between the subunits, supporting a hypothesis that these ribosomes are highly translationally active. While for this system with only two species, the cell biology and architecture are different enough within a lamella to determine whether a feature of interest is microbe- or host-derived, this ability to differentiate between subtomogram averages may be important in systems with more species or, for example, in future work to establish vesicle origin in the crypt space.

### Bacteria inside the crypt spaces

The cryo-ET and acquired FS-ET data confirmed the previous TEM findings, which state that the bacteria are in close proximity to the microvilli of the host tissue. However, the data revealed morphological complexity in the crypt space, with some areas having a very narrow space of single-file bacteria (Fig. 5), particularly when the bacteria are releasing symbionts into the environment as they do each dawn ^32^. The bacteria within the crypt space were not tightly packed but rather interspaced with crypt space lumen as well as many vesicles. Interestingly, previous resin-embedded samples, as well as the data obtained here through FS-ET and blockface SEM, have shown a seemingly denser packing of bacteria within the crypts than in cryo-ET, possibly stemming from dehydration artefacts during resin embedding or freeze substitution.

**FIG 5.**
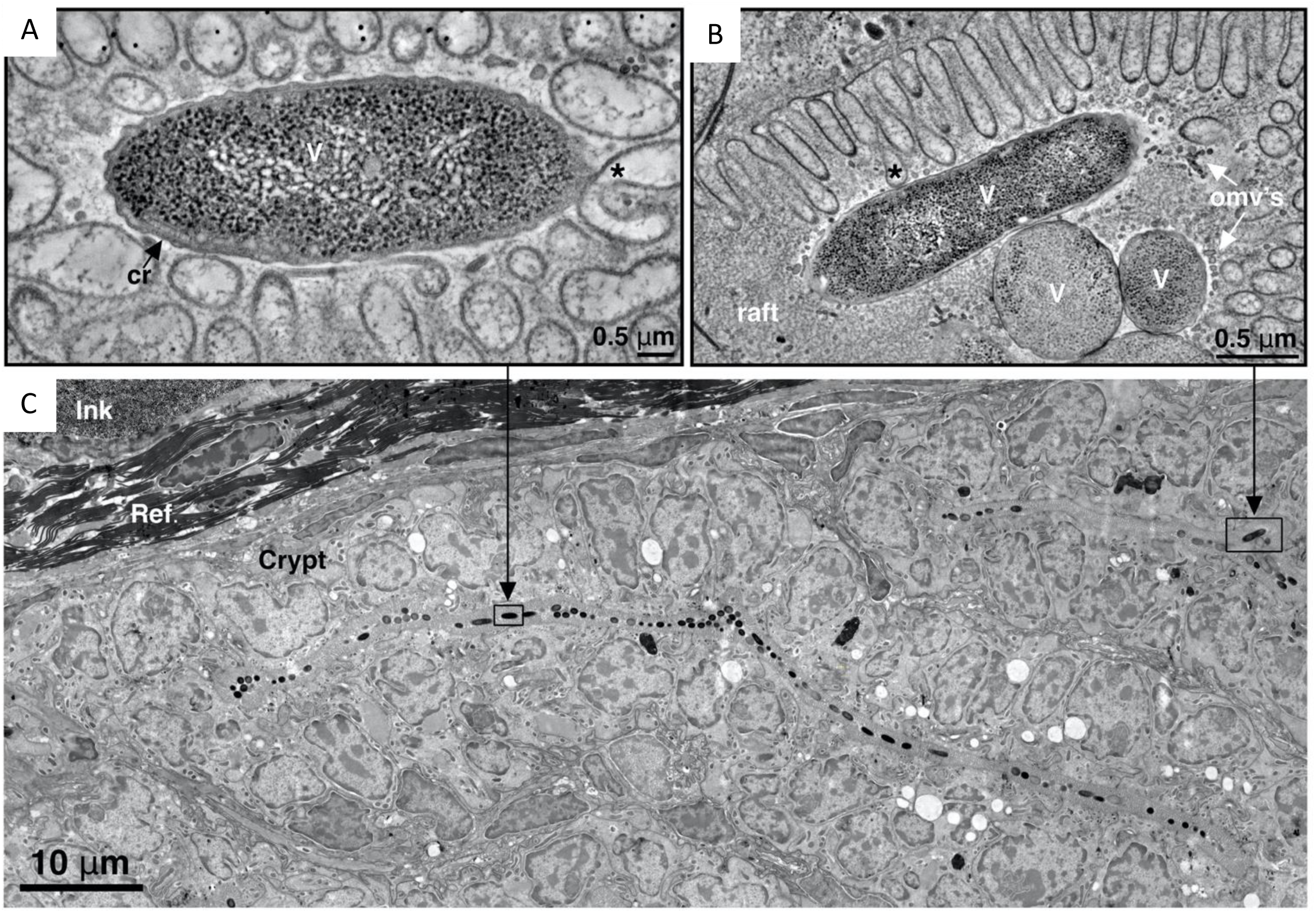
Freeze substituted electron tomography of symbiotic squid tissue. (A-B) Individual *V. fischeri* surrounded by microvilli in crypt space. (C) Biogeography of host tissue surrounding crypt space showing narrow line of symbionts. Shown: (cr) chemoreceptor, (*) host microvilli (Ref) reflector (c), ink sac (is, ink), reflector (r, ref). Scalebar: noted above.

Within the bacterial cells and in their proximity, we observed a variety of bacterial nanostructures, including intracellular filament bundles of unknown identity (Fig. 3F), chemoreceptors (Fig. 3G), type-6 secretion systems (T6SS) (Fig. 3I), flagella, and ribosomes (Fig. 3G-I). It is believed that *V. fischeri* relies on secretion systems, T6SS1 and T6SS2^33^, to regulate symbiosis and bacteria-bacteria interactions before and during symbiosis. We show a discharged T6SS2 structure within the crypt space where there does not appear to be a nearby active target (Fig. 3I).

We also discovered a phage within the crypt space (Fig. 3H). The capsid head is empty, supporting the likelihood that the phage has already injected its DNA into its target. The phage head and sheath are near bacteria in the tomogram, with the missing wedge effect masking whether the phage is still docked to the bacteria situated above it in the tomogram.

### Symbiont communication

Progressing to the host-bacterial interface, the main question we asked is related to information traffic. To better understand this dynamic, we focused on two key ways of communication: host-bacterial contact and vesicular traffic. We found that the bacteria are localized near the microvilli that line the boundaries of the crypt space (Fig. 6A-B). While not all bacteria were in contact with microvilli, we observed 18 instances where they were in close proximity to microvilli. For these instances, the distance between the host membrane and bacterial outer membrane was 13.7 ± 3.1 nm (Fig. 6C). This intermembrane gap was generally visibly denser, i.e. darker, than the general crypt space lumen, indicative of biological material spanning the distance between the two species. Additionally, at certain contact points, the bacterial outer membrane would adopt a concave curvature to accommodate the microvillus tip (Fig. 6A).

**FIG 6.**
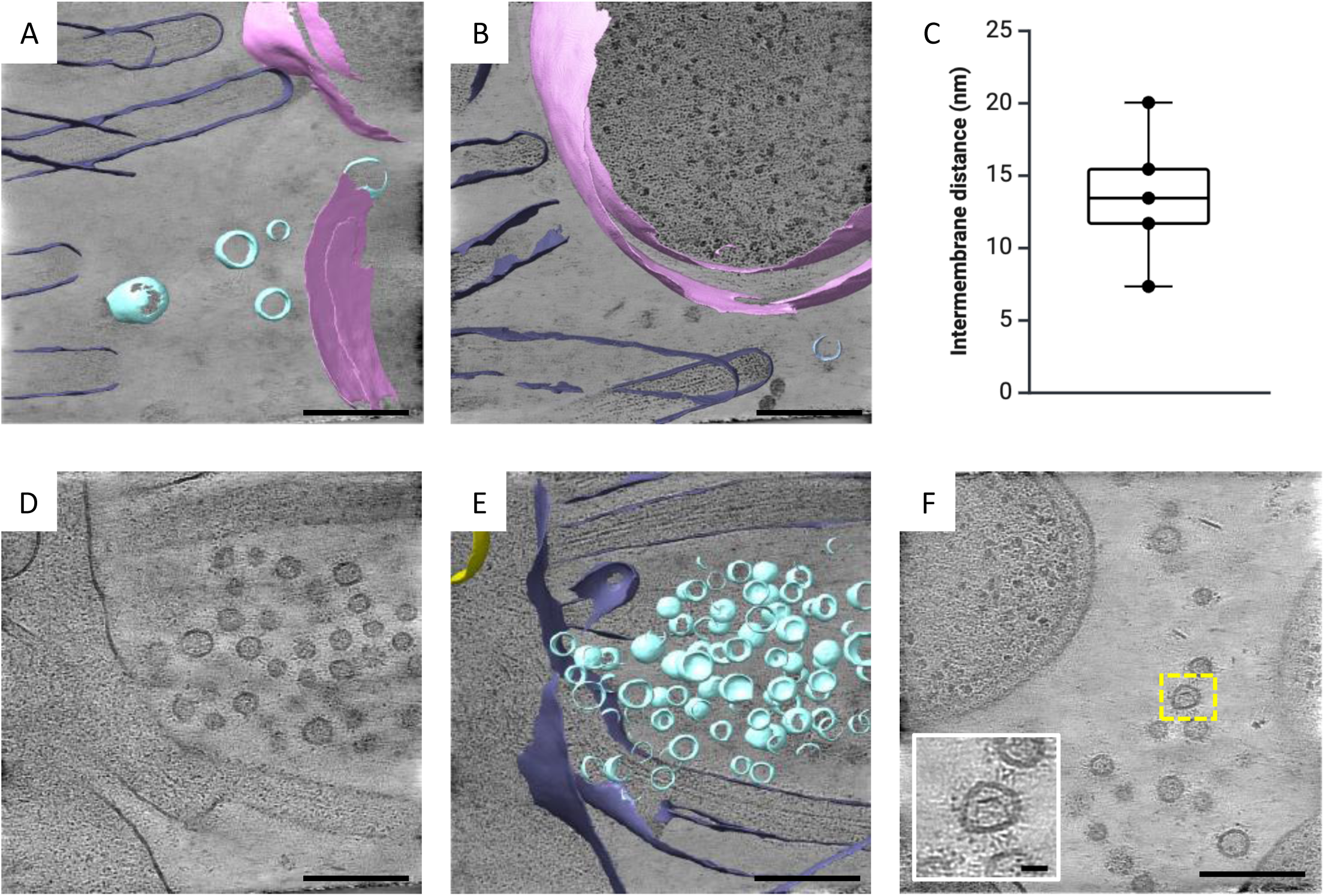
CryoET of host-microbe interface. (A-B) Tomograms and segmentation showing host-microbe interface. Host microvilli (purple), *V. fischeri* symbiont (pink), vesicles (blue). (C) Plot of intermembrane distance between host microvilli and bacterial outer membranes (13.7 ± 3.1 nm). (D-E) Tomogram and segmentation showing a burst of vesicles at the intersection between two epithelial cells. Vesicles (blue), host epithelial membrane (purple). (F) Host-derived, vesicle encapsulated, flotillin-like structures (yellow box) found within the crypt space. Scalebars: (A-B, D-F) 200 nm, (F) inset 20 nm.

We also explored strategies alternative to direct contact. Previous studies have shown that outer membrane vesicles (OMVs) are secreted from bacteria to be taken up by the host ^24,25^. The assumption was therefore that most vesicles present within the crypt space would be bacteria-derived. While this idea may still be true, there are many vesicles within the crypt of various sizes and coating, that may be host-tissue derived. These include vesicles within the crypt space containing flotillin-like structures (Fig. 6F). While flotillins are known to exist through all branches of biology, the only ones to have been solved structurally, and which resemble those found within these vesicles, are from mammalian sources. The likelihood that these vesicles are host-tissue derived is further supported by the fact that we observe these flotillin-like structures throughout the animal tissue, including the apical membranes of the crypt space epithelia (Fig. 3C). Another indication that a number of vesicles found in the crypt space are of host origin is the cluster of vesicles often found at the junction between two epithelial cells (Fig. 6D-E). While, in theory, there could be a mechanism by which bacterial-derived OMVs are clustered at these positions, it seems more likely that vesicles are by some means released in bursts at these interfaces. These vesicles are interconnected by density which perhaps results in the coatings of some of the vesicles seen throughout the crypt space. Additionally, it is expected that host microvilli-derived vesicles might also be present in the crypt spaces (see e.g., ^21^).

## Discussion

While our understanding of host-microbe systems in terms of the impact of the symbionts on the animal transcriptome is rapidly growing^9,34,35^, their well-defined interactions at the structural level are still largely unknown. Here we aimed to gain such insight using one of the simplest known natural host-microbe pairs: the symbiosis between the bioluminescent bacterium *Vibrio fischeri* and the Hawaiian bobtail squid, *Euprymna scolopes*. We applied a multi-modal imaging pipeline to process complex native tissues for cryo-ET. Each step in the imaging processing pipeline gains resolving power, but at the loss of contextual information. This approach allowed us, for the first time, to investigate the interaction between host tissue and microbes in a near native state, at macromolecular resolution while maintaining the greater volumetric context of the Squid-Vibrio system.

Our current understanding of the host crypt space has been largely driven by conventional TEM and confocal microscopy (for review^7^). Our methods here confirm the historical understanding of the symbiotic landscape. Bacteria-bacteria and bacteria-host spacing and interaction was also represented in our tomograms without the added artefacts of fixation, dehydration, or heavy-metal staining. Existing imaging methods have limited the capability of discretely measuring those distances and fully interpreting the mechanisms for interactions at the host-microbe interface. Here, we could precisely quantify the distance between symbiont and the host’s microvilli. Indeed, this new lens at the interface brings up fundamental questions regarding the physical processes and mechanisms of communication which could occur over distances of ∼13.4 nm.

Within the Squid-Vibrio symbiosis, type-6 secretion systems have been implicated in regulating bacterial populations and driving gene expression in host tissues^22,23,33,36^ Complex interactions of bacteria within culture and within animal tissue have been recorded previously using light microscopy. Our data reveal for the first time a discharged T6SS2 within crypt space (Fig. 3I). While not visibly in contact with a target at the time of vitrification, it could be that the contact site was outside of the ∼200 nm lamella height. However, we were able to confirm the presence and use of T6SS within colonized crypt spaces.

We were also able to demonstrate how subtomogram averaging can be used to differentiate between host and symbiont macromolecules (Fig. 4 G, H). Additionally, it was used to explore molecules that have never been previously determined *in situ*, such as haemocyanin (Fig. 4D-E) and even used to tentatively determine the identity of macromolecules through their structure, i.e. squid acetyl CoA-carboxylase (Fig. 4A-C). Taken together these examples exemplify the potential of STA to explore complex, symbiotic systems.

While this work here clearly demonstrates that cryo-ET can be used as an invaluable tool to gain insight into the structural interaction of a host with microbiome, we still face some challenges. First and foremost, sample preparation and vitrification still remain technically challenging. High-pressure freezing remains the enigma within the cryo-ET pipeline, as the technology for freezing has been relatively stagnant for decades. There is currently no reliable way to quantify or qualify vitrification levels of samples or deep within samples until the lamella is imaged in the TEM, which is expensive and time consuming. We believe coordinated efforts to redesign vitrification methods for very large samples needs to be prioritized, with innovation in vitrification checks early in the sample pipeline. Initial sample processing had low success rates, but by the publishing of this paper, the majority of HPF samples were suitable for FIB-milling, dependent on accessibility of region of interest for lift out region.

In addition to vitrification, we need to optimize existing correlative cryo-fluorescent imaging for deep tissues which will increase accuracy during targeting. Every step taken to ensure proper vitrification and accurate targeting early on will save time and resources in later imaging and processing steps. Accurate lateral targeting within the sample is generally achievable with cryo-confocal imaging. However, with large volume, 10-200 µm deep, samples, the axial dimension is still limiting. Serial lift-out does help with this challenge as it allows data collection at multiple depths in the tissue in the serial sections and, in turn, increases the chance that the exact region of interest is included in the final data set. In the future, however, a combination of coarse removal of material via microtomy or FIB-milling and then acquiring fluorescence of the newly generated cliff-edge, should result in depth resolution by imaging the depth of the sample with the lateral resolution of the light microscope^37^. This approach should result in an improved and more consistent targeting.

Now that we have demonstrated the feasibility of this approach, future plans are to investigate the host-microbe interface in more detail with hosts colonized by different strains of symbiont, and at different time points during the initiation and maintenance periods of symbiosis. These different *Vibrio fischeri* strains have differing colonization abilities, and result in different host responses where remote tissues of the squid sense and respond to symbiotic state and the diel rhythm for presence of bacteria ^8,9,38,39^. Ongoing efforts are to expand imaging to investigate other squid tissue, including the skin, eyes, gills, tentacles, and accessory nidamental gland.

The Squid-Vibrio system is a model organism for host-microbe interface for reasons including its binary nature, bypassing the need for microbe identification at the tomography level. However, many other symbiotic tissues and host-microbe associated tissues have relationships with a multitude of microbial partners, which are currently difficult or impossible to differentiate at the tomography level. Current work is underway to develop methods for microbial differentiation labeling at the TEM level.

## Methods

### Colonization Protocol

Juvenile *E. scolopes* were hatched from clutches in artificial seawater (ASW) at Leiden University, Leiden, The Netherlands. Hatchling squid were colonized overnight with GFP-labeled *V. fischeri* strain ES401 (ES401-GFP) using methods previously described^40^. Colonized animals were confirmed for downstream processing by the presence of symbiont luminescence detected by a luminometer (Turner Designs, Inc., Sunnyvale, CA, USA).

### Sample preparation for Micro-CT and plastic-section electron microscopy

Squid light organs were dissected from anesthetized squid in 2% EtOH in ASW and fixed with 3% glutaraldehyde, 1% paraformaldehyde, 5% sucrose in marine PBS. Samples were rinsed with fresh marine PBS containing 10% Ficoll, placed into brass planchettes (Ted Pella, Inc.), and rapidly frozen with a Wohlwend Compact-03 high-pressure freezing machine (Technotrade International). The vitrified samples were transferred under liquid nitrogen to cryotubes (Nunc) containing a frozen solution of 2.5% osmium tetroxide, 0.05% uranyl acetate in acetone. Tubes were loaded into an AFS-2 freeze-substitution machine (Leica Microsystems, Vienna) and processed at −90°C for 72 h, warmed over 12 h to −20°C, held at that temperature for 6 h, then warmed to 4°C for 1 h. The fixative was removed, and the samples rinsed 4x with cold acetone, after which they were infiltrated with Epon-Araldite resin (Electron Microscopy Sciences) over 48 h. Samples were flat-embedded between two Teflon-coated glass microscope slides and the resin polymerized at 60°C for 48 h.

Embedded samples were observed by light microscopy to ascertain sample quality and orientation, then excised from the slide with a scalpel and remounted onto plastic sectioning stubs with a 2-part epoxy adhesive. Serial semi-thick (170 nm) sections were cut with a UC6 ultramicrotome (Leica Microsystems, Vienna) using a diamond knife (Diatome, Ltd. Switzerland). Sections were collected onto formvar-coated 1 mm slot copper-rhodium grids (Electron Microscopy Sciences) and stained with 3% uranyl acetate and Reynold’s lead citrate. For some experiments gold beads (10 nm) were placed on both surfaces of the grid to serve as fiducial markers for subsequent tomographic image alignment.

### Micro-CT

Epon-embedded light organs were stabilized with beeswax on an Xradia holder for MicroCT data collection. The samples were scanned with an Xradia Versa 520 (Zeiss Xradia 520 Versa, Carl Zeiss., CA, USA). Scout-and-ScanTM Control System was used to set the scan setting. Material was scanned using 80 KV, 2 seconds exposure time, and 4x objective resulting in a voxel size of 1.88 µm. Scout-and-Scan Control System Reconstructor was used for the MicroCT data reconstruction. The images were analysed and manipulated using Avizo software (Version: 8.01; Thermo, Fisher Scientific).

### Electron microscopy and Dual-Axis tomography

Grids were placed in a dual-axis tomography holder (Model 2040, E.A. Fischione Instruments) and imaged with a Tecnai T12-G2 transmission electron microscope operating at 120 KeV (Thermo Fisher Scientific) equipped with a 2k x 2k CCD camera (XP1000; Gatan, Inc.). Tomographic tilt-series and large-area montaged overviews were acquired automatically using the SerialEM software package ^41^. Automated electron microscope tomography using robust prediction of specimen movements ^41^. For tomography, samples were tilted ± 62° and images collected at 1° intervals. The grid was then rotated 90° and a similar series taken about the orthogonal axis. Tomographic data was calculated, analyzed, and modeled using the IMOD software package ^42,43^. Correction for non-perpendicularity of beam and tilt Axis in tomographic reconstructions with the IMOD package ^43^. Automated tilt series alignment and tomographic reconstruction in IMOD ^44^) on iMac Pro and Mac Studio M1 computers (Apple, Inc.).

### Serial BlockFace-SEM

Samples were prepared for block face SEM according to a slightly adapted protocol from ^45^. After fixing the material for 2 h at room temperature with 2.5% glutaraldehyde + 2% paraformaldehyde in 0.15 M cacodylate buffer containing 2 mM CaCl2, the material was washed 3 times with cacodylate buffer and then placed into 2% OsO4 / 1.5% potassium ferrocyanide in 0.15 M cacodylate buffer containing 2 mM CaCl2. The material was left for 60 minutes on ice. After washing 3 times in milliQ water, the material was placed into 1% Thiocarbohydrazide for 20 minutes at room temperature. The material was again washed with milliQ water and then stained with 2% aqueous OsO4 for 30 min at room temperature. After washing 3 times with milliQ water, the material was placed into 1% Uranyl acetate for 2 hours at room temperature. The material was washed with milliQ water then stained with Lead aspartate for 30 minutes at 60 °C. The material was washed with milliQ water and then dehydrated on ice in 20%, 50% and 70% ethanol solutions for 5 minutes at each step. After replacing the 70% ethanol with a fresh 70% ethanol solution, the samples were kept overnight at 4°C. The next day, samples were dehydrated in 90%, 100%, 100% ethanol solutions for 5 minutes at each step. Next, the material was kept in dry acetone for 10 minutes on ice, and another 10 minutes in fresh dry acetone at room temperature. The material was infiltrated with 25%, 50% and 75% Spurr (Sigma-Aldrich) solution in acetone for 2 hours at room temperature each step, followed by an overnight step at room temperature in 100% Spurr resin (Sigma-Aldrich). The next day, the material was placed in fresh Spurr resin for 2 hours at room temperature, after which the material was embedded and polymerized at 60 °C for 48 hours.

Data was collected with a Gatan 3View2XP unit installed on a Zeiss Gemini 300 field emission SEM. Fourteen volumes containing between 173 to 846 sections were collected at 1.6 kV accelerating voltage and focal charge compensator at 65%. The pixel dwell time was three microseconds, the pixel size was 10 or 15 nm and the section thickness of 100 nm.

### High pressure freezing, fluorescent targeting and Serial Lift-Out of vitrified tissue

In order to allow for subsequent targeting of the crypt areas by fluorescence, colonized hatchlings were stained for 30 mins using CellTracker^TM^ Orange CMRA (Molecular Probes, Invitrogen). The hatchlings were rinsed twice and then anesthetized in 2% EtOH in artificial seawater (v/v) for 5 mins. The light organs were then dissected and frozen either using a Bal-Tec HPM 010 (Bal-Tec) or a Leica EM ICE (Leica Microsystems). Three mm type A and type B carriers (Leica Microsystems) were used, and coated in 1-hexadecene, removing excess liquid with filter paper (Whatmann). The light organs were placed into the 200 µm well of the type A carriers filled with 20% Ficoll 400 in ASW (w/v) such that the ink sac faced down and the reflector up. The flat side of a type B carrier was place on top, sealing the sample, and the carrier sandwich was then high pressure frozen. Samples were stored in LN2 until further use.

To accurately target the host-microbe interface during Serial Lift-Out, samples were then imaged using cryo-fluorescence. The HPF samples were loaded into a shuttle designed and manufactured in-house which, in turn, was loaded into a Leica TCS SP8 laser confocal microscope (Leica Microsystems), equipped with a cryo-stage. The grids were imaged using the LAS X software (3.5.5.19976, Leica Microsystems), a 50×, 0.9 NA objective, and using HyD and PMT detectors. To enable proper targeting, three channels were acquired: GFP, CellTracker^TM^ Orange CMRA, and reflection, labeling the *V. fischeri*, the host tissue and the sample surface, respectively. The GFP signal was acquired with an λ_ex_ = 488 nm at 10% total laser power and λ_em_ = 496-530 nm. The CellTracker^TM^ Orange signal was acquired with λ_ex_ = 552 nm at 1% total laser power and λ_em_ = 560-609 nm. Reflection acquired with an λ_ex_ = 552 nm at 1% total laser power and λ_em_ = 549-555 nm. In order to acquire data over the depth of the the sample, tile set montages were acquired of the entire carrier. with a pinhole size of 6.39 AU and a voxel size of 0.578 µm × 0.578 µm × 5.001µm. The resulting image stacks were merged and the maximum-intensity projections were determined using LAS X.

Serial Lift-Out was performed in accordance to the protocol published in Schiøtz, *et al.* and for milling patterns and finer details one is referred there ^5^. In order to ensure proper targeting, trench milling was performed on a METEOR (Delmic) equipped Aquilos 1 FIB-SEM system (Thermo Fisher Scientific). The HPF sample was loaded into a 45° cryo-FIB-SEM shuttle and then into the microscope. Once loaded, the METEOR was used to confirm the crypt region, previously localized with the cryo-confocal. This data was acquired using Odemis version 3.2.1, with a LMPLFLN 50×/NA = 0.5 objective. The GFP signal was acquired with an λ_ex_ = 484 nm and an λ_em_ = 525/30 nm. The CellTracker^TM^ Orange CMRA was acquired with an λ_ex_ = 600 nm and an λ_em_ = 607/36 nm. The reflection signal was acquired with an λ_ex_ = 625 nm and no emission filter. Once localized, the METEOR-FIB transposition was used to define the milling area for trench milling. Before this, however, the sample was coated in a protective layer of organometallic platinum for 90 s with the stage at a pre-defined position (WD = 10.6 mm, stage tilt = 45. Trench milling was then performed with the sample perpendicular to the ion beam (trench-milling orientation) using a mill-and-check approach, regularly checking the trench milling progress with the METEOR, in order to confirm proper targeting. After three side of the block had been created the stage was rotated 180° (lamella-milling orientation), and tilted to 25° to perform the undercut, releasing the bottom of the extraction block from the bulk HPF material. The resulting block was ∼50 µm × 40 µm, with a depth from ∼20-30 µm.

The lift-out and sectioning was performed by loading the trench-milled HPF carrier and a 100/400 mesh copper grid in an Aquilos 2 (Thermo Fisher Scientific) equipped with an EasyLift micromanipulator (Thermo Fisher Scientific). The previously milled trench was centered with the stage in trench-milling orientation. The copper block adapted needle micromanipulator was then inserted into the FIB chamber and brought in contact with the upper surface of the extraction volume. An array redeposition milling at 1 nA was used to attached the biology to the needle. The side of the block still remaining was then ablated and the volume was extracted.

Double-sided attachment Serial Lift-Out was chosen as the attachment method, due to its improved stability. The shuttle was brought into lamella-milling orientation and tilted to 18° and the micromanipulator was reinserted into the chamber. The extraction volume was then maneuvered between the 400 mesh grid bars, removing excess material from the side of the block if needed. An array of redeposition milling done at 1 nA on each of the grid bars welded the biology to the grid. The needle was then maneuvered 50-100 nm up and a 1 nA line pattern placed 2 µm above the edge of the volume, as well as two line patterns along the blocks left and right edges, were used to release the section. The remaining volume was moved to a new lift-in location and the process was iterated until the block was depleted.

The receiver grid with the 20 sections was transferred to an Aquilos 1 FIB-SEM (Thermo Fisher Scientific) equipped with a METEOR (Delmic). The sections were then screened for the presence of *V. fischeri* using the integrated light microscope. The GFP signal was acquired with an λ_ex_ = 484 nm and an λ_em_ = 525/30 nm. The reflection signal was acquired with an λ_ex_ = 625 nm and no emission filter. The resulting data was used to guide the position of the milling of the ∼20 µm wide lamella within the section.

Lamella milling was performed stepwise, to 1.5 µm, 800 nm and 500 nm, at 0.5 nA, 0.3 nA and 0.1 nA, respectively. Lamellae were then fine milled to their final at 50 pA to a final thickness of ∼200 nm. Finally, over- and under-tilting by 0.5° was used to even the lamella thickness. The resulting lamella were sputter coated twice with metallic platinum for 4 s (current = 15.0 mA, chamber pressure 0.20 mbar).

### Cryo-electron tomography data acquisition

Tilt series were collected on a Selectris X energy filter and Falcon 4i equipped 300 kV Titan Krios G4 instrument (Thermo Fisher Scientific). Data was collected using Tomo5, version 5.12.0 (Thermo Fisher Scientific). An altas of the grid was acquired by at magnification of ×125 and lamella montages were acquired at a magnification of ×11,500 (2.127 nm pixel size).

Tilt series were acquired at a magnification of ×64,000 (1.89 Å pixel size), over a tilt range of −70° to 50°. The Hagen scheme was used to acquire the data in a dose-symmetric manner, using a tilt increment of 2°, and a dose of 2e^−^/Å^2^ per tilt (total dose = 120 e^−^/Å^2^). The defocus values were cycle at 0.5 µm increments between −1.5 and −5 µm.

### Tomogram reconstruction and subtomogram averaging of haemocyanin, ribosome

#### Tomogram reconstruction

Tomograms were reconstructed using cryoBOOST^215^, an in-house wrapper program which runs RELION 5.0^46^ as well as performing automated removal of bad tilts. MotioncCor2 version 1.4.0^47^ was used for motion correction, grouping 29 EERs per frame, as well as splitting the dataset into half sets required for subsequent denoising. CTFFIND4 version 4.1.14^48^ was used for CTF estimation and AreTomo version 1.3.3^49^ was used for tilt series alignment. RELION 5.0 was used to reconstruct the 3D volume and Cryo-CARE was used for denoising, with resulting binned and denoised tomograms having a pixel size of 11.72 Å.

#### Filament subtomogram averaging

Initial particle picking was performed on five tomograms containing the crystalline-like arrays by tracing the filaments using AMIRA version 2021.2^50^ (Thermo Fisher Scientific). This was adapted from a protocol using cylinder correlation described by Schneider *et al.* These segmentations were then used to oversample the filaments, with a sampling distance of 30 Å^51^.

The 96,850 positions were then extracted at a pixel size of 7.56 Å (bin4) and a box size of 128 pixels and used to generate an initial average using RELION 5.0^46^. The particle positions were then refined, limiting sampling by applying priors for the tilts to 15°. The resulting particle list was then distance cleaned to remove duplicates caused by the oversampling. This was done by clustering the positions generated during the alignment. This reduced the list down to 13228 particles.

Particles were re-extracted at a pixel size of 1.89 Å (bin1) and averaged. A mask was then generated using a threshold of 0.0013, extended by 3 pixels and applying a 15-pixel soft edge. The particles were then processed by 3D refinement, with a prior of 15° tilt, resulting in a ∼21 Å structure. This was then submitted to three iterations of Bayesian polishing^52^, CTF refinement and 3D refinement. This was finally post processed, producing a 14.2 Å density map, GSFSC.

#### Hemocyanin subtomogram averaging

Initial particle picking was performed manually on a subselection of tomograms using the ChimeraX version 1.6.1^53^ plugin ArtiaX^54^. 414 particles were extracted and used to create an initial structure at pixel size 7.56 Å (bin4) using RELION 5.0^46^. This structure was used as a template for a subsequent iteration of template matching using PYTOM version 0.7.1 on all 10 tomograms containing regions of blood sinus. Using RELION 5.0, 10,108 particles were extracted, aligned and classified at a pixel size of 15.12 Å (bin8), using C1 symmetry. Bad particles were removed and further processing was done with D5 symmetry applied as the outer wall of the particle displayed D5 symmetry, confirming what would be expected from previous studies^31^. After further alignments and classifications, a total of 9999 particles were retained. These were submitted to multiple iterations of alignment, CTF refinement and Bayesian polishing at a pixel size 1.89 Å (bin1). The resulting D5 symmetrical structure showed an average resolution of 9.9 Å, GSFSC.

#### *V. fischeri* ribosome subtomogram averaging

Template matching was initially performed on a few tomograms using a reference of a 20 nm sphere and was done using PYTOM version 0.7.1^55,56^. Using RELION version 5.0^46^, 1196 particles were extracted and used to create an initial structure with a pixel size of 7.56 Å (bin4). This structure was then used as a template for a subsequent iteration of template matching using PYTOM version 0.7.1. This was performed on all tomograms of the dataset containing *V. fischeri*. Using RELION version 5.0,13,594 particles were extracted, aligned and classified at a pixel size of 15.12 Å (bin8). Poorly aligned particles were removed and the remaining 8,434 particles were reextracted at a pixel size of 1.89 Å (bin1). These were subjected to focused refinement of the small and large ribosomal subunits, using masks generated with the Segger tool in ChimeraX version 1.6.1^53^. The structures averaged to a resolution of 9.7 Å and 8.7 Å, GSFSC, for the small and large subunit, respectively. ChimeraX version 1.6.1 was used to dock these into a previously obtained lower resolution full ribosome structure.

#### *E. scolopes* ribosome subtomogram averaging

Template matching was initially performed on a few tomograms using a reference of a 30 nm sphere and was done using PYTOM version 0.7.1^55,56^. Using RELION version 5.0^46^, 1591 particles were extracted and used to create an initial structure with a pixel size of 7.56 Å (bin 4). This structure was then used as a template for a subsequent iteration of template matching using PYTOM version 0.7.1. This was performed on all tomograms of containing *E. scolopes* tissue. Using RELION version 5.0, 30,503 particles were extracted, aligned and classified at pixel size 15.12 Å (bin8). Poorly aligned particles were removed and the remaining 15,883 particles were reextracted at a pixel size of 1.89 Å (bin1). Multiple iterations of alignment, CTF refinement and Bayesian polishing were performed, resulting in a structure at a resolution of 7.5 Å, GSFSC.

#### Segmentation and distance analysis

Tomograms were segmented using MemBrainSeg version 0.0.5^57^, using the pre-trained neural network available. Segmentation analysis was done in IMOD version 4.12.32^42,44^, with distances being determined by measuring the closest distance between the segmentations of the microvilli and *V. fischeri* membranes. The 18 measurements were then averaged to determine the mean distance and standard deviation (13.7 ± 3.1 nm). Distance at the mitochondrial-plasma membrane close contact was determined without prior segmentation by distance measurement in IMOD version 4.12.32^42,44^.

## Acknowledgements

This work was supported by funding from the Gordon and Betty Moore Foundation (Symbiosis Model Systems, Grant #9328). This work was partly funded by the Max Planck Society and the European Union’s Horizon Europe Projects IMAGINE (GA#101094250) and ‘SymPore’ (951292). This study used the infrastructure of the Department of Cell and Virus Structure at the MPI of Biochemistry. This work was also partly funded by the M. J. M. Murdock Charitable Trust (#1-339-02548). The authors thank Vera Williams for the illustration in Figure 2.

---

**Supplementary Movie 1.**
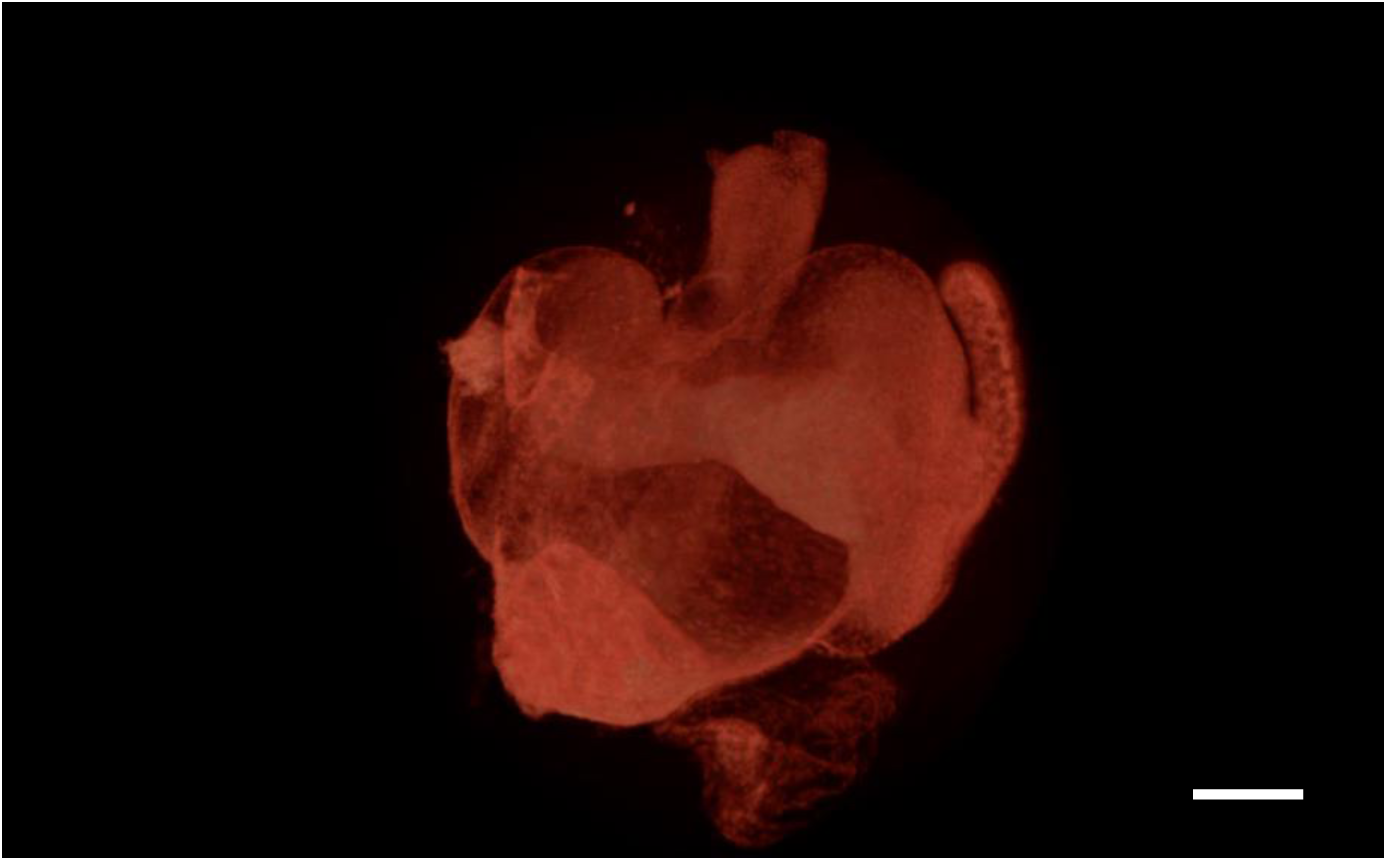
Micro-CT of juvenile *E. scolopes* light organ. Scalebar: 50 µm.

**Supplementary FIG 1.**
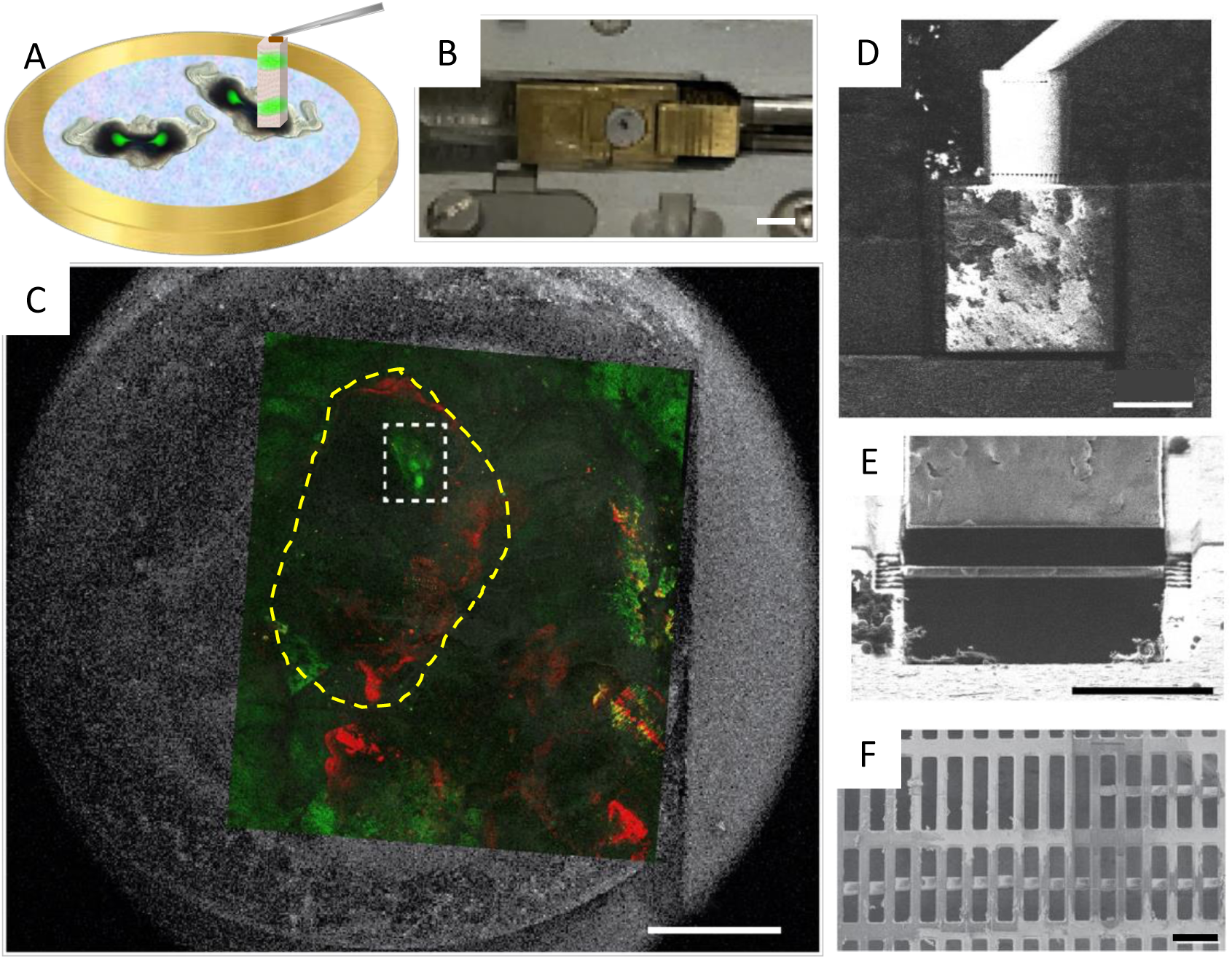
HPF and FIB-SEM Lift-out of vitrified, colonized squid tissue. (A) Colonized squid light organ is HPF in planchet and (B) loaded into FIB-SEM directly in HPF planchet. (C) Cryo-fluorescent correlative image is overlaid onto sample, yellow dashed region outlines one light organ, white dashed box is one crypt space. Lift-out targeted the crypt space for maximum host-microbe interface (GFP*: V. fischeri*, RFP: CellTracker, host epithelium). (D-F) Cryo-lift-out of host-microbe interface. Lift-out block is removed from HPF planchet, and deposited onto lift-in grid by serial sectioning using FIB, and attached to both sides of grid bars of 400x100 grid. Sections are thinned into lamella about 200 nm thick. Scalebars: (B) 3mm (C) 500 µm (D) 20 µm (E) 20 µm (F) 100 µm.

**Supplementary FIG 2.**
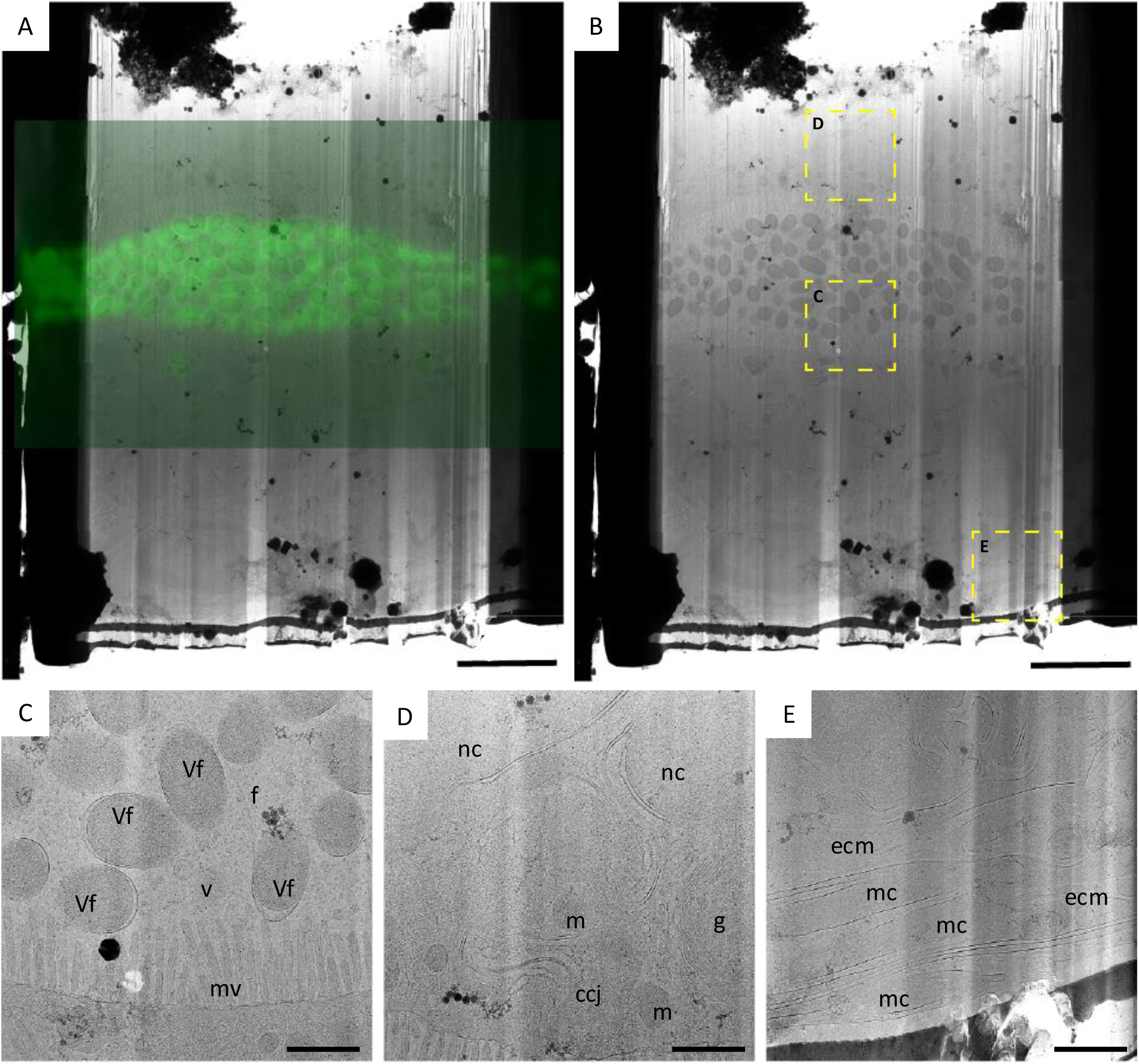
Correlation and overview of host-microbe interface in vitrified lamella. (A) Fluorescent signal of GFP-labeled *V. fischeri* in crypt space correlated with a lamella overview. (B) A lamella overview, with regions of interest outlined in yellow, with corresponding cell cell biology in (C-E) Labels: *Vibrio fischeri* (Vf), vesicle (v), microvilli (mv), nucleus (nc), mitochondria (m), Golgi apparatus (g), cell-cell junction (ccj), extracellular matrix (ecm), muscle cell (mc). Scalebars: (A-B) 5 µm, (C-E) 1 µm.

**Supplementary FIG 3.**
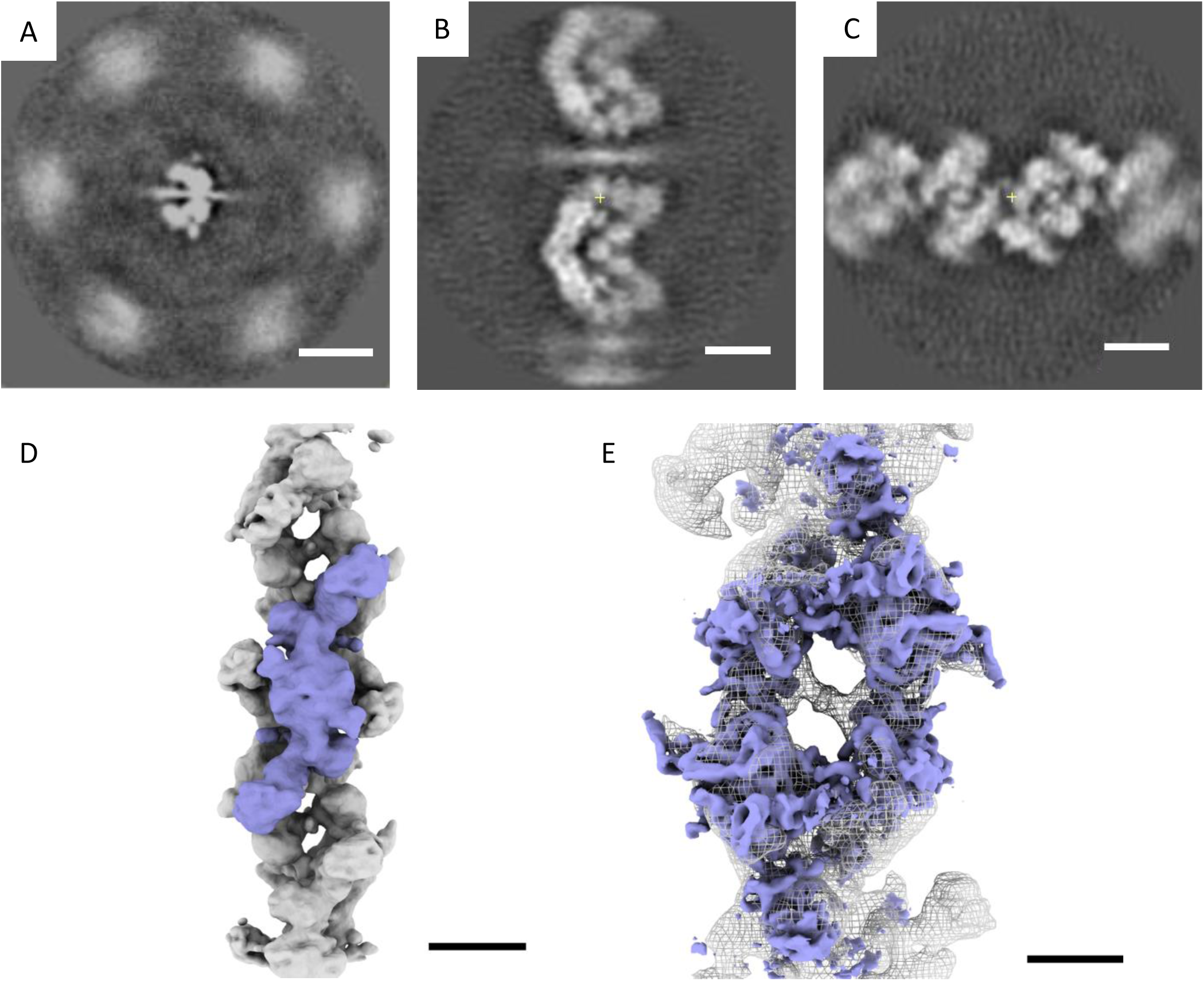
Subtomogram average of squid crystalline-like filaments. (A) Top view of the subtomogram filament average, depicting the hexagonal packing, as the beginning of a coiled-coil protrusion extending out from the central filament to the neighboring ones. (B, C) Side-views of the same filament. In (B) the coiled-coil extensions are also clearly visible. (D) Structure of the filament averaged to a resolution of 14.2 Å, GSFSC. Purple indicates the homodimer, three of which make up the ∼290 nm helical rise of the filament. (E) The docking of the inactive state of human acetyl-CoA carboxylase (PDB ID 6G2H, purple) within the filament density (grey). The density shows a large degree of similarity, with the exception of the coiled-coil domain, which may explain the crystalline-like array, not previously reported in human^30^. Scalebars: (A) 20 nm, (B-D) 10 nm, (E) 5 nm.

**Supplementary FIG 4.**
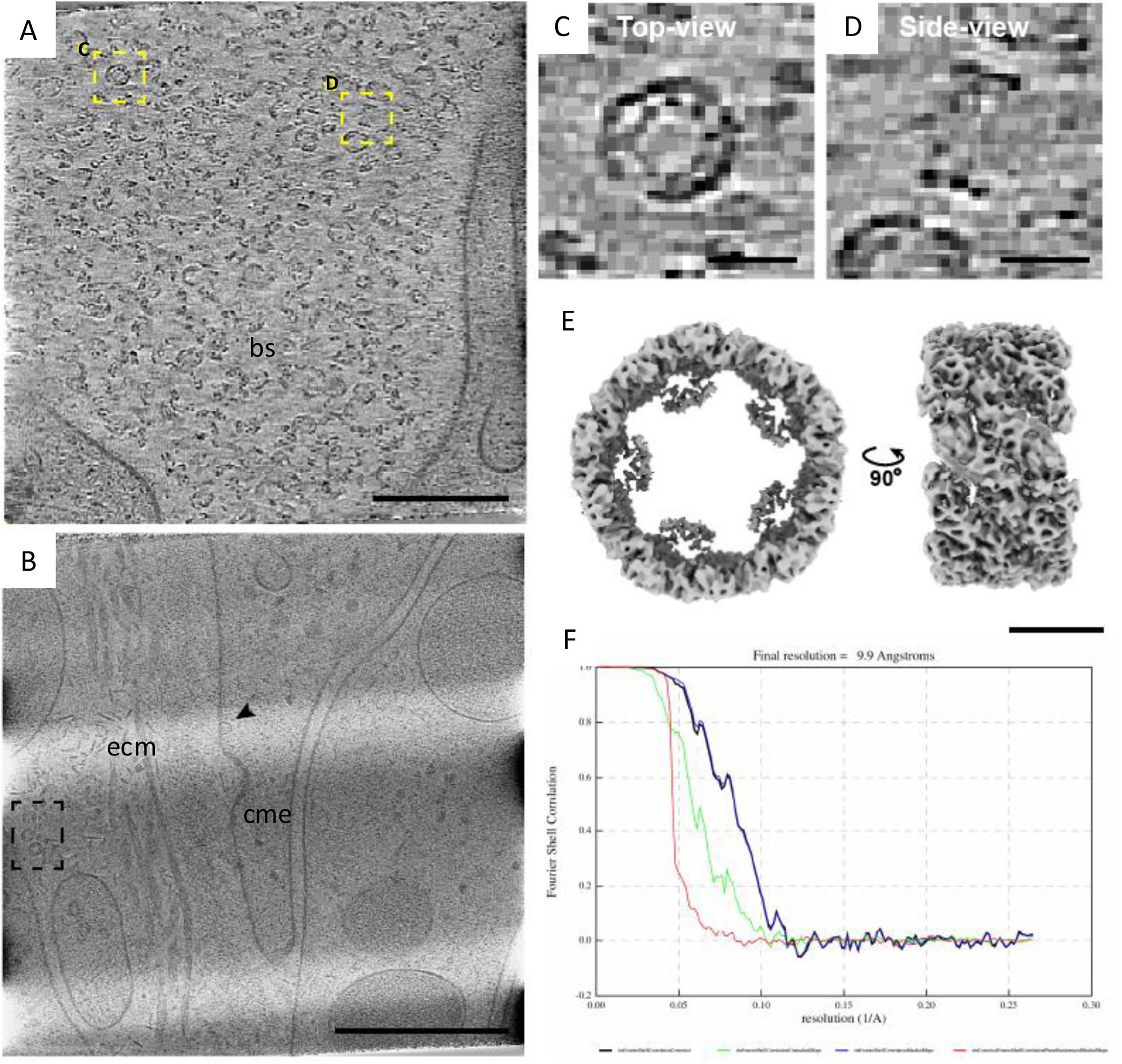
CryoET and subvolume average of haemocyanin in host tissue. (A) Blood sinus (bs) of squid tissue filled with haemocyanin proteins, individual haemocyanin marked with yellow-dashed box (B) Haemocyanin (black dashed box) near extracellular matrix (ecm) during clatherin-mediated endocytosis (cme), black arrow marks flotillin structure. (C-D) Top and side view of haemocyanin structure. (E) Subtomogram average of haemocyanin, with D5 symmetry applied. (F) The FSC plot of the structure, depicting that the GSFSC extends to 9.9 Å resolution. Scalebars: (A) 200 nm (B) 400 nm (C-D) 200 Å (E) 10 nm.

**Supplementary FIG 5.**
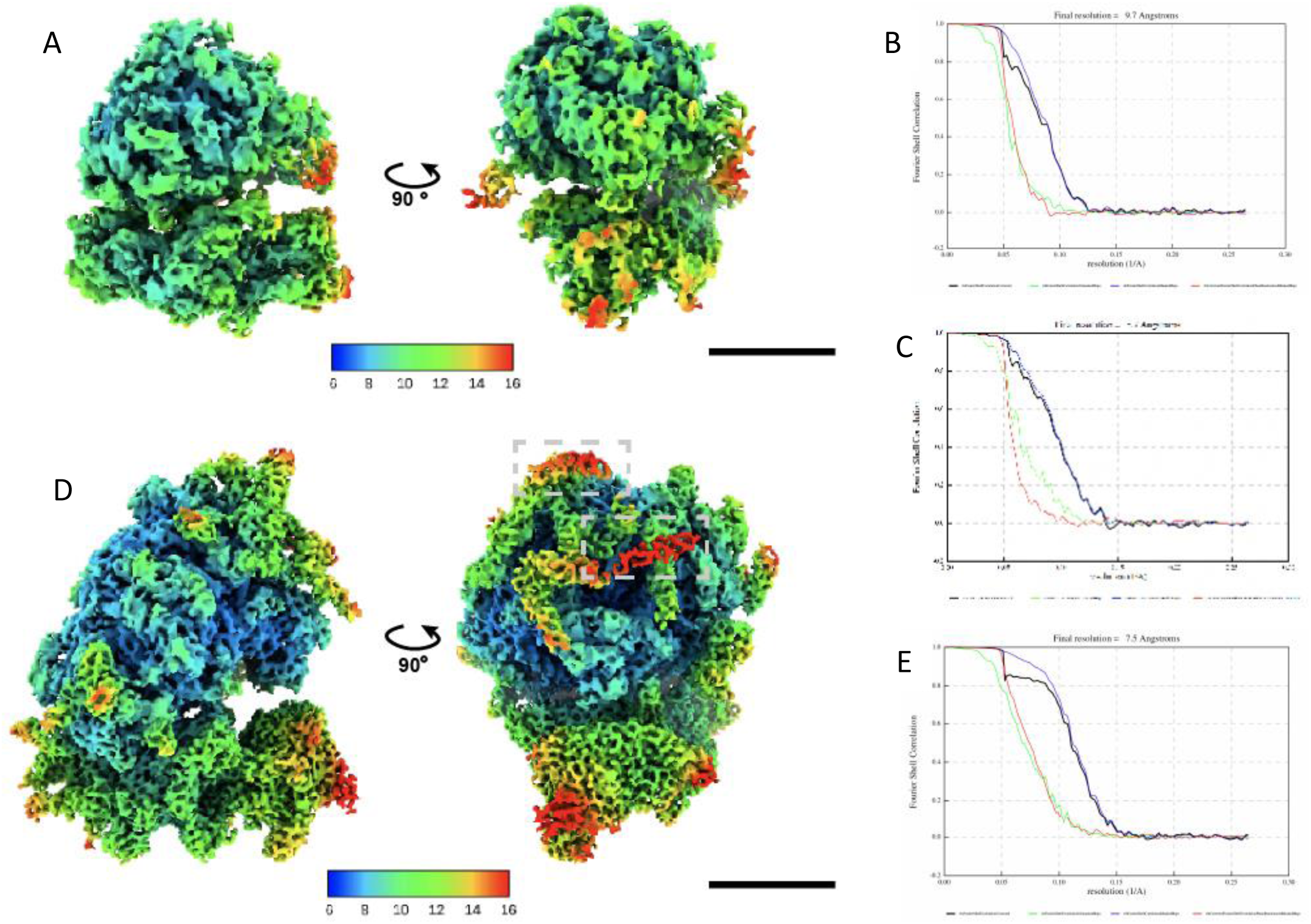
Subtomogram averages of squid ribosome and *V. fischeri* ribosome. (A) Subtomogram average of the *V. fischeri* ribosome. Color coating depicts local zresolution. (B, C) FSC curves for both the small and large subunit of the *V. fischeri* ribosome, depicting that they resolve to 9.7 Å and 8.7 Å, respectively (D) Subtomogram average of the *E. scolopes* ribosome. Color coating depicts local resolution (E) FSC curve for the *E. scolopes* ribosome, depicting that it resolves to an average resolution of 7.5 Å. Scalebars: (A) 10 nm (D) 10 nm.

